# Comprehensive transcriptomic and proteomic profiling of mTOR-related epilepsy

**DOI:** 10.64898/2026.03.10.710809

**Authors:** Kaiyuan Ji, Satoshi Miyashita, Keiya Iijima, Kaoru Yagita, Tomoo Owa, Kazumi Shimaoka, Nao K N Tabe, Minami Mizuno, Ayane Hosaka, Kayo Nishitani, Kumiko Murayama, Kanako Komatsu, Masaki Sone, Terunori Sano, Shinichiro Taya, Tomoki Nishioka, Kozo Kaibuchi, Masaki Takao, Masaki Iwasaki, Mikio Hoshino

## Abstract

Malformations of cortical development (MCDs) are a major cause of drug-resistant epilepsy (DRE) in children. A subset of MCDs, including focal cortical dysplasia type II, tuberous sclerosis complex, and hemimegalencephaly, shares characteristic histopathological features and somatic mutations in genes within the PI3K–AKT–mTOR signaling cascade, and is termed mTORopathies. However, the molecular mechanisms that drive epileptogenesis and consequent uncontrollable seizures in these disorders remain poorly understood.

Here, we performed integrated genomic, transcriptomic, and proteomic analyses of 60 surgical brain specimens from 42 patients with MCDs and 18 with mesial temporal lobe epilepsy.

Targeted genomic sequencing identified 19 somatic mutations in PI3K–AKT–mTOR pathway genes, including novel variants in *AKT3* (L51H) and *MTOR* (A512T). RNA sequencing comparing mTORopathy brains with controls, along with cross-validation using an independent public dataset, identified 287 consistently differentially expressed genes that were consistently altered. Downregulated genes were enriched in oxidative phosphorylation (OXPHOS) and mitochondrial metabolic pathways, whereas upregulated genes were associated with gliogenesis and cellular senescence. Proteomic profiling identified 44 differentially expressed proteins, including significant reductions in OXPHOS-related proteins (COX5B, NDUFS4, GOT2, RAB27B, and DKK3).

Our study provides a patient-matched, multi-omics dataset of human mTOR-related epilepsies from a single-institution cohort. Integrated analyses across transcriptomic and proteomic layers highlight impairment of OXPHOS as a molecular hallmark of mTORopathies, Integrated transcriptomic and proteomic analyses identified impaired OXPHOS as a molecular hallmark, potentially contributing to focal hypometabolism in mTORopathy. These findings offer mechanistic insights into epileptogenesis and provide a valuable resource for understanding molecular pathologies and developing mechanism-based therapies for mTOR-related epilepsy.

## Introduction

Epilepsy is one of the most common neurological disorders, affecting approximately 50 million people worldwide, and is characterized by recurrent, unprovoked seizures.^1^ While nearly 70% of individuals with epilepsy can control their seizures with antiepileptic drugs, the remaining 30% suffer from drug-resistant epilepsy (DRE), where seizures remain uncontrollable despite medication.^2^

Malformations of cortical development (MCDs) are a group of rare neurodevelopmental disorders, characterized by abnormalities in brain size and/or morphology, caused by neurodevelopmental defects such as neurogenesis or neuronal migration.^3^ MCDs are a leading cause of DRE in pediatric patients (∼40%), and are associated with severe neurological consequences, including lifelong epilepsy, intellectual disability, and behavioral abnormalities, placing a substantial burden on patients worldwide.^4^ A specific subgroup of MCDs—including focal cortical dysplasia type IIa/b (FCD IIa/b), tuberous sclerosis complex (TSC), and hemimegalencephaly (HME)—shares distinctive histopathological features such as dysmorphic neurons, enlarged glial cells, and aberrant activation of the mechanistic target of rapamycin (mTOR) signaling pathway.^5,6^ Recent genomic studies of brain tissues from patients with FCD IIa/b, TSC, and HME identified somatic mutations in genes involved in the PI3K/AKT/mTOR signaling cascade.^7–11^

Supporting these findings, mouse models with constitutive mTOR pathway activation in the cerebral cortex exhibit migration defects, enlarged cells, and spontaneous recurrent seizures.^9,12,13^ Together, these studies indicate that somatic mutations and subsequent mTOR hyperactivation represent a key trigger for epilepsy in this subset of MCDs. As a result, these disorders are currently referred to as mTOR-related epilepsies or “mTORopathies”. However, the cellular and molecular mechanisms underlying epileptogenesis and drug-resistant seizures in mTORopathy patients remain elusive.

Integrated transcriptomic and proteomic analyses provide a more comprehensive and accurate understanding of the molecular landscape of neurological disorders. In this study, we conducted a comprehensive transcriptomic and proteomic characterization of resected brain specimens obtained from Japanese patients with mTORopathies and stored at a single institute, in order to elucidate the molecular mechanisms driving epileptogenesis and DRE. Our integrated analysis of transcriptomic and proteomic data confirmed the activation of previously reported pathways involved with gliogenesis and cellular senescence. In addition, we identified a significant reduction in oxidative phosphorylation signals (OXPHOS), providing molecular insight for focal hypometabolism observed by FDG-PET^14,15^ and implicating mitochondrial energy metabolism as a potential contributor to epileptogenesis. Our comprehensive datasets represent a valuable resource for understanding the molecular basis of epilepsy in mTORopathies and identifying potential molecular targets for future treatments.

## Materials and methods

### Patient selection

In this study, we included 56 patients who (1) underwent neurosurgical resection for drug-resistant focal epilepsy at the National Center Hospital, National Center of Neurology and Psychiatry (NCNP), Kodaira, Tokyo, Japan; (2) were diagnosed with focal cortical dysplasia (FCD), tuberous sclerosis complex (TSC), or hemimegalencephaly (HME) by experienced histopathologists; and (3) had sufficient high-quality RNA and frozen tissue available for transcriptomic and proteomic analyses among the 60 patients from 2002 to 2021. As a control group, we used temporal tip cortex specimens obtained from patients with mesial temporal lobe epilepsy associated with hippocampal sclerosis who underwent neurosurgery at NCNP. Control specimens were confirmed to be neurotypical by histopathological assessment. Clinical information and specimens were obtained according to the Declaration of Helsinki with written informed consent. This study was approved by the NCNP Ethics Committee, Japan (NCNP-A2018-050). Primary outcomes for this report were (i) differential gene expression profiles from bulk RNA sequencing and (ii) differential protein abundance profiles from LC–MS/MS proteomics, comparing mTORopathy lesions with control cortex.

### Genome analysis

Genomic DNA was extracted from frozen brain tissue using the QIAamp DNA Micro Kit (QIAGEN) according to the manufacturer’s instructions. Sequence libraries were constructed using the Ion AmpliSeq Library Kit Plus (Thermo Fisher Scientific), and the concentration was measured with the Agilent High Sensitivity DNA Kit (Agilent) on the Agilent 2100 bioanalyzer (Agilent).

A 50pM sequence library was loaded on an Ion Chef™ with the Ion 540 Kit Chef and Ion 540 Chip (Thermo Fisher Scientific) to prepare sequence template. Sequencing was carried out on the Ion GeneStudio S5 System (Thermo Fisher Scientific). Sequence data were analyzed using Ion Reporter Software (Thermo Fisher Scientific) with the human reference genome hg19. Multiple primer sets covering exonic and exon-intron border regions (+25 to −25) of target genes involved in PI3K/AKT/mTOR signaling (Table S) were designed using the Ion AmpliSeq Designer software (Thermo Fisher Scientific).

### Histopathological diagnosis

Brain tissues were fixed using 4% paraformaldehyde in 0.1 M phosphate buffer (pH 7.4) and paraffin embedded. Some parts of the samples were stored at -80°. All samples were routinely stained with H&E and KB staining. Immunohistochemistry was performed by using anti-NeuN (A60 mouse monoclonal, cat# MAB377, Millipore, USA; 1:500), and anti-neurofilament H (SMI-32 mouse monoclonal, cat# SMI-32R, Covance, USA; 1:4,000) antibodies. Immunohistochemical staining was automatically performed by the VENTANA system (Roche, Basel, Switzerland) according to the manufacturer’s instructions. All histopathological diagnoses were assessed by well-experienced neuropathologists.

### RNA extraction and RNA sequencing

RNA was extracted from epileptogenic lesions from malformations of cortical dysplasia and temporal lobe epilepsy with hippocampal sclerosis, which were surgically resected and cryopreserved in the NCNP Biobank. RNA-sequencing libraries were created using SMART-seq and Nextera XT DNA Library Prep kit. Sequencing libraries were sequenced on a NovaSeq 6000.

### Processing of RNA sequencing data

Quality filtering of sequenced fastq files was performed using fastp (v.0.20.0). Filtered sequences were then aligned with STAR (v. 2.7.10) to the GRCh38 human reference genome. Aligned reads were then counted using htseq (v.0.11.3) with GTF.

Identification of differential expression genes (DEGs) 41 MCD patients in this cohort were compiled as the MCD group. MCD patients with FCD IIa, FCD IIb, TSC, and HME were assigned to the mTORopathy group; MCD patients with FCD I and HS with FCD (FCD III) were assigned to the non-mTORopathy group. Using the DESeq2 R package, genes with adjusted P < 0.05 as differentially expressed genes (DEGs).

### GO Enrichment Analysis

Gene ontology enrichment analysis was performed by using the “enrichR” package in R. DEGs generated from whole MCD samples, the mTORopathy group, the non-mTORopathy group, and each specific MCD type were used as the lists of DEGs for GO analysis. Adjusted *P* value < 0.05 was identified as a cut-off for statistical significance in the enrichment results.

### Gene Set Enrichment Analysis

Molecular Signatures Database accessed by the “msigdbr” R package provides thousands of annotated gene sets for use with fast gene set enrichment analysis via the “fgsea” R package.

Hallmark gene sets (with argument: category = "H") consist of various well-defined biological states or pathways.

### Data acquisition and processing of proteomics data

Frozen brain sections were powdered by using cyro-press CP-100WP (Microtech Nition) according to the manufacturer’s instructions. Powdered samples were lysed with 50mM Tris-HCl, 8M urea, 0.005% bromophenol blue, 1% SDS and sonicated for 20 sec. Protein concentration of each sample was measured by BCA assay (Fujifilm). 5ug proteins were purified and digested by SP3 protocol (Hughes et al., 2019). Protein was reduced with 10 mM dithiothreitol at RT for 30 min and alkylated with 20 mM iodoacetamide at RT for 30 min in the dark. Samples were mixed with 150 μg of magnet beads (a 1:1 mixture of hydrophilic and hydrophobic SeraMag carboxylate-modified beads). Ethanol was mixed to a final concentration of 50%. The samples were mixed by a ThermoMixer C (Eppendorf, Hamburg, Germany) at 1200 rpm for 5 minutes at 24°C. The protein-bound beads were washed 3 times with 80% ethanol by using a magnetic rack. Beads were resuspended with 100uL of 50 mM ammonium bicarbonate. Trypsin/Lys-C Mix (1:50, w/w) was added to the sample tubes. Protein digestion was performed at 37°C with continuous mixing at 1,300 rpm overnight in a ThermoMixer C. Digested peptides were collected and desalted by using GL-Tip-SDB and resuspended in 20μL of 2% acetonitrile, 0.1% TFA solution. Liquid chromatography–mass spectrometry(LC/MS) was performed on a Vanquish Neo UHPLC system (Thermo Scientific) coupled with Orbitrap Fusion (Thermo Scientific). Peptide samples were loaded onto a trap column (Acclaim PepMap 100 C18 LC column, 3 μm, 75 μm ID × 20 mm; Thermo Fisher Scientific) and were separated on an EASY-Spray C18 LC column (75 µm × 15 cm, 3 µm, 100 Å, Thermo Fisher Scientific) by a linear gradient consisting of 0%–35% acetonitrile with 0.1% formic acid over 120 min at a flow rate of 300 nl/min.

In the data-independent acquisition (DIA) method, MS1 scan range was set at 390–1010 m/z in positive mode, orbitrap resolution at 120,000, and standard AGC target mode with maximal injection time of 60 ms. MS2 scan range was set 400–1001 m/z with orbitrap resolution at 60,000 and normalized HCD collision energy was set at 30%. Number of scan event was 50 with 12 m/z isolation windows in quadrupole mode.DIA data was processed using DIA-NN (v1.9.2).

Default settings for in-silico spectral library preparation were used. A human protein database from UniprotKB (UP000005640_2024_10_18.fasta) was used to prepare an in-silico spectral library. DIA-NN analysis was performed using default settings (Trypsin/P digestion, one missed cleavage is allowed, carbamidomethylation on cysteine as fixed modification, acetylation on the N-term as variable modification, false discovery rate for peptides and proteins were set to 0.01, match between runs was enabled).

Proteins containing any missing values across samples were removed prior to downstream analyses. Differential protein expression analysis between the Control and mTORopathy groups was assessed using linear modeling with empirical Bayes moderation implemented in the limma package with the Benjamini–Hochberg method to obtain adjusted *P* values. To compare the expression levels of COX5B, NDUFS4, GOT2, DKK3, and RAB27B among the control, FCD I, FCD III, and mTORopathy groups, we performed ANOVA followed by Tukey’s post hoc test for multiple comparisons.

### Public Data acquisition

The expression profiles and corresponding clinical information of MCDs patients derived from one public dataset (GSE256068) were downloaded from GEO database (https://www.ncbi.nlm.nih.gov/geo/). Then we used the same R packages for the subsequent analysis including DEGs identification, GO terms enrichment analysis, and fast gene set enrichment analysis.

## Results

### The clinical features and genomic analysis of MCD patients

We analyzed 56 surgical brain specimens obtained from 56 patients who underwent neurosurgical resection at the National Center of Neurology and Psychiatry (NCNP). Among them, 41 samples were derived from patients diagnosed with focal cortical dysplasia type I or II (FCD I/II), tuberous sclerosis complex (TSC), or hemimegalencephaly (HME), while 15 samples of lateral temporal neocortex were obtained from patients with mesial temporal lobe epilepsy (mTLE) caused by hippocampal sclerosis (Fig. 1A, Table S1). 10 tissues without cortical dysplasia were regarded as the control group (Fig. 1B), and five tissues with focal cortical dysplasia were regarded as FCDIIIa (HS+FCD). 41 MCD patients included in this study consisted of 24 males and 17 females (Table S1). Among them, 10 samples were obtained from temporal lobe (TL), 26 samples were from frontal lobe (FL), three sample from parietal lobe (PL), one sample from occipital lobe (OL), and one sample across TL and FL. Since all MCD and control samples were derived from the Japanese ethnic group, the effect of genetic heterogeneity was considered to be minimal. The average age at surgery was 5.4 years old for MCD patients and 34.3 years old for controls.

**Figure 1.**
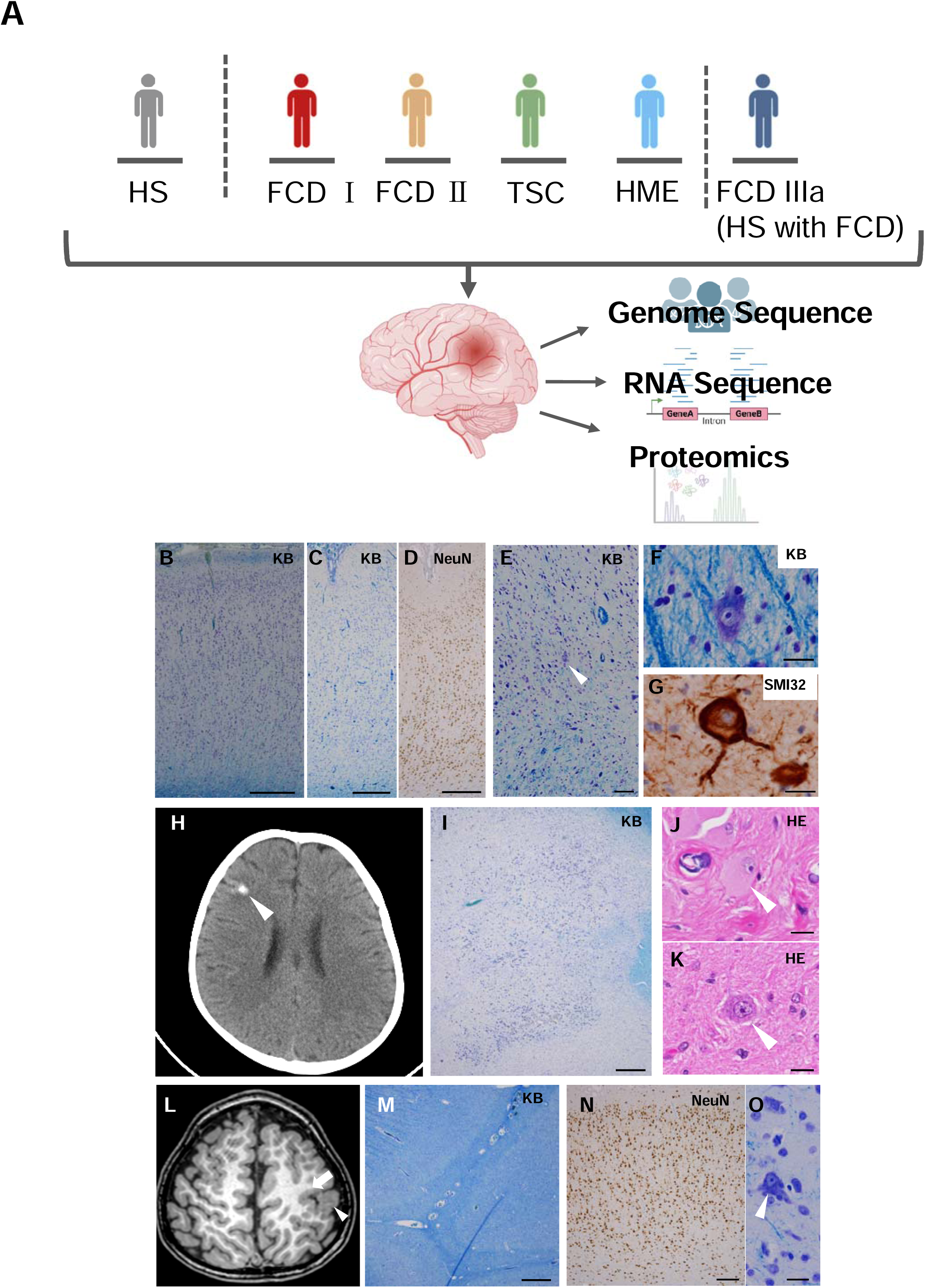
Overview of experimental design and neuropathological features of MCD patients. (A) Schematic illustration of patient groups included in the study: controls (HS), and malformations of cortical development (MCD), including FCD type I, FCD type IIa/b, tuberous sclerosis complex (TSC), hemimegalencephaly (HME), and HS with cortical disruption (FCD III). Genome sequencing, RNA sequencing, and proteomics analysis were performed. (B) A photomicrograph of the lateral temporal cortex from a control patient. Klüver-Barrera (KB) stain. (C,D) Photomicrographs of the cerebral cortex of FCD type 1 show disorganized cortical layers. KB stain (C). Immunostaining of NeuN (D). (E-G) Photomicrographs of the cerebral cortex of FCD type IIa show disorganized cortical lamination (E) and dysmorphic neurons (E arrow, F, G). KB stain (E, F). Immunostaining of SMI-32 (G). (H-K) A brain CT image of a TSC individual shows high-intensity areas consistent with a cortical tuber (H, arrowhead). Photomicrographs of the cortical tuber show disorganized layer structure and accumulation of neurons (I), a balloon cell (J), and a dysmorphic neuron (K). KB stain (K) and HE stain (J, K) (L–O) A brain MRI image shows the megalencephalic region of HME (L, arrowhead). Photomicrographs of the cerebral cortex of HME individual show dyslamination of neurons (M, N), and a dysmorphic neuron (O). KB stain (M, O). NeuN immunohistochemistry (N). Scale bars = 500 μm (B-D, I, M-N), 100 μm (E), 20 μm (F-G, J-K, O)

Typical histopathological features of FCD I (layer dyslamination, Fig. 1C, D), FCD II a/b (presence of dysmorphic neurons and balloon cells, Fig. 1 E-G), TSC (tuber structure and abnormal cells, dysmorphic neurons and giant cells, Fig. 1H-K), and HME (megalencephaly in the affected hemisphere with abnormal cells, Fig. 1L-O) were visualized by histological and imaging analyses.

We first performed genomic sequencing targeting PI3K/AKT/mTOR signaling pathway-related genes by using the genomic DNA extracted from resected lesions (Table S2). We found 19 somatic mutations in the genomic region of six mTOR-related genes (*MTOR, TSC2, AKT3, AKT1S1, PIK3CA, and PIK3R2)* from the surgically resected lesions of 18 patients (Table S2, mutation rates 2.56% to 35.95%). Eight of the somatic mutations were pathogenic mutations previously reported to increase the mTOR activity. Two somatic mutations in *TSC2* and *AKT1S1,* known suppressors of mTOR signaling, were frame shift mutations, potentially leading to the activation of mTOR signaling. Of note, we detected novel mutations in *AKT3* gene (AKT3 L51H) and *MTOR* gene (MTOR A512T) from HME and FCD II patients.

### Global transcriptomic characteristics of MCD brains

To investigate the molecular features underlying the epileptogenesis and/or uncontrollable seizures in mTORopathy, we performed RNA sequencing (RNA-seq) analysis using epileptogenic lesions resected from MCD brains, including FCDIIa/b, HME, and TSC. Temporal lobe regions of the neocortex resected from temporal lobe epilepsy patients with hippocampal sclerosis were used as a control (Table S1). To assess the global transcriptomic features of MCDs, FCD III, and control, the hierarchical clustering was performed using genes with counts greater than five across the entire dataset (Fig. 2A). Nine controls were clearly clustered and almost separately distributed from MCD and FCD III samples. Age at surgery and sex of patients did not profoundly affect the transcriptional profiles (Fig. 2A). In addition, there were no obvious clusters within the MCD subtypes, suggesting that transcriptional profiles of the epileptic lesions in MCDs share common transcriptional features compared to those in controls.

**Figure 2.**
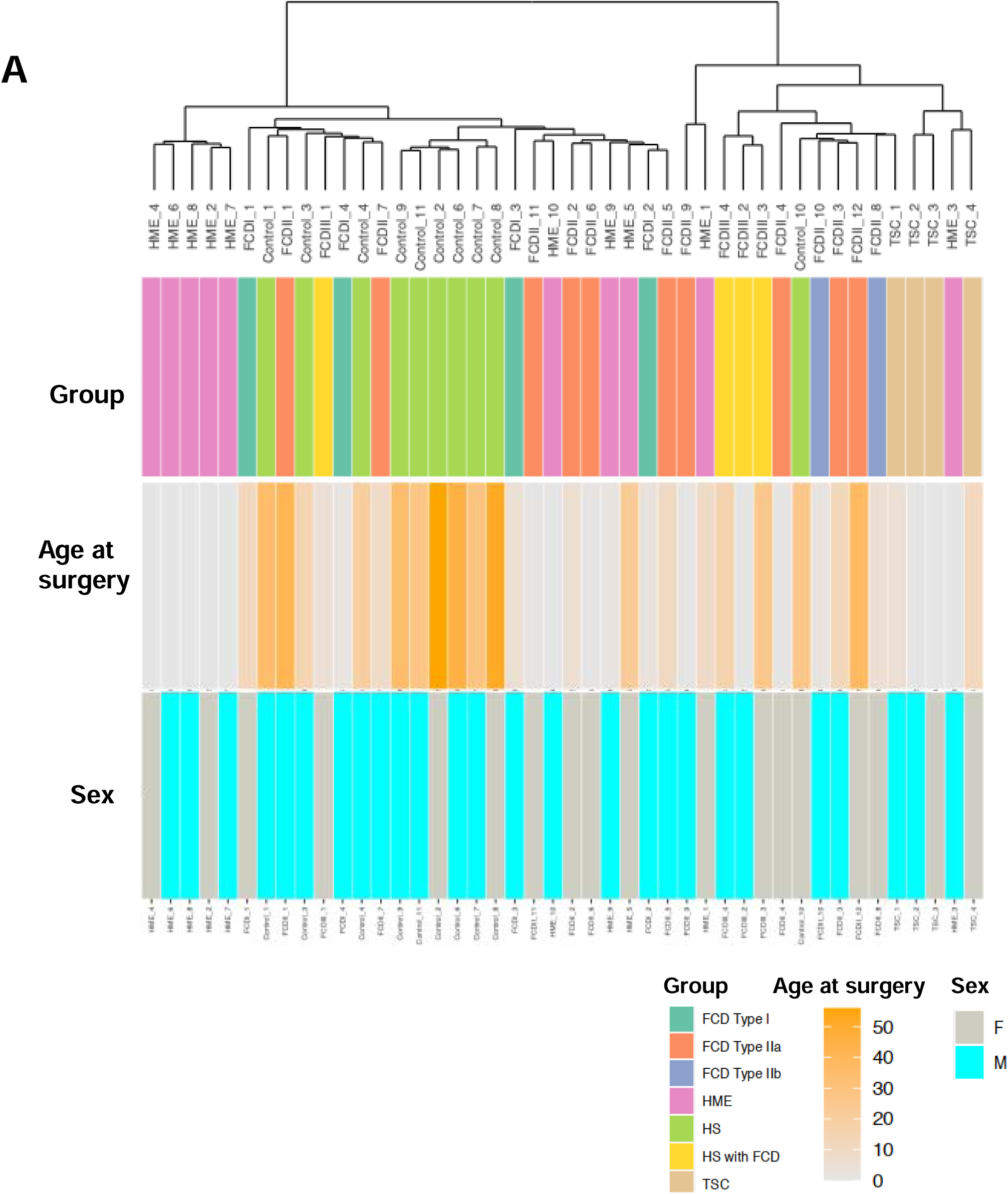
Patient cohort characteristics. (A) Hierarchical clustering of transcriptomic profiles of all samples based on transcriptomic expressions. Clinical metadata shown below: group classification, age at surgery, and sex.

### Alterations of gene expression in mTORopathy brains

Differential expression analysis between the control and mTORopathy groups revealed that expressions of 650 genes were significantly increased while 395 genes were decreased in the mTORopathy group (Fig. 3A, Table S3,4). Next, we compared DEGs from our own dataset with previously published RNA-seq dataset^16^ to further confirm whether the identified DEGs in this study were common molecular features of mTORopathy. From the François et al. dataset, we extracted 45 patients with mTORopathy (27 FCDIIa/b and 18 TSC) and 12 control data, and identified a total of 6,605 DEGs between the control and mTORopathy groups (3,024 upregulated, 3,583 downregulated in the mTORopathy group. By comparing the DEGs between our and the François et al. dataset^16^, 115 genes were identified as commonly increased genes in mTORopathy, and 52 genes were as commonly decreased genes (Fig. 3B,C. Table S5). These 167 commonly dysregulated genes might represent key molecular mediators underlying epileptogenesis and drug-resistant seizures in the mTORopathies.

**Figure 3.**
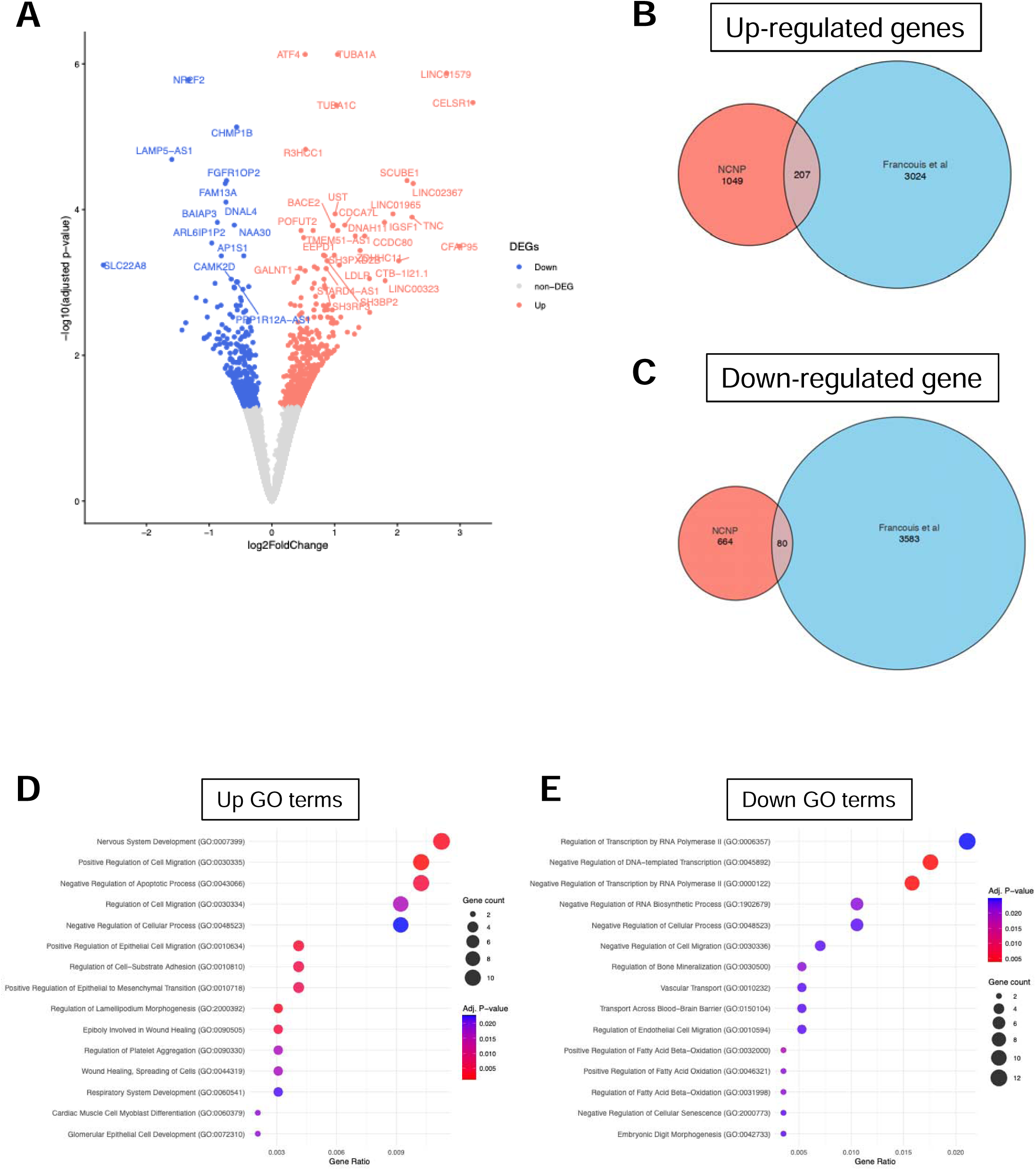
Differential expression analysis between mTORopathy and controls. (A) Volcano plot showing significantly upregulated (red) and downregulated (blue) genes (DEGs). Selected transcripts are labeled. (B, C) Venn diagrams comparing upregulated (B) and downregulated (C) genes identified in this study (NCNP cohort) with previously published data (François et al.). (D) Gene Ontology (GO) terms enriched among upregulated genes. (E) GO terms enriched among downregulated genes.

### Characterization of altered molecular and signaling pathways in mTORopathy

To explore the biological functions of commonly dysregulated genes (Table S5), we next performed gene ontology (GO) enrichment analysis with GO terms of biological process (BP) terms associated with the commonly upregulated 207 genes and downregulated 80 genes, respectively. 30 GO terms related to upregulated genes (up GO) and 6 GO terms related to downregulated genes (down GO) were detected as significant (Fig. 3D, E. Table S6). Among GO terms related to the upregulated genes, apoptosis-related GO terms including “Negative Regulation of Apoptotic Process (GO:0043066)”, and “Regulation of Apoptotic Process (GO:0042981)” were identified. In addition, neural development-related GO terms, such as “Nervous System Development (GO:0007399)”, “Regulation of Cell Migration (GO:0030334)”, and “Negative Regulation of Stem Cell Differentiation (GO:2000737)” were detected, possibly reflecting the defects in neural development in mTORopathy brains. Among the GO terms related to the downregulated DEGs, fatty acid metabolism-related GO terms including “Positive Regulation of Fatty Acid Beta-Oxidation (GO:0032000)”, “Positive Regulation of Fatty Acid Oxidation (GO:0046321)”, and “Positive Regulation of Lipid Catabolic Process (GO:0050996)” were identified with the tissue development-related GO terms including “Negative Regulation of Cell Migration (GO:0030336)”, “Negative Regulation of Cell Differentiation (GO:0045596)”, and “Regulation of Cell Migration (GO:0030334)”.

To estimate dysregulated signaling pathways, we performed gene set enrichment analysis (GSEA) using hallmark gene sets from the Human Molecular Signatures Database. GSEA was conducted separately on our dataset and the public dataset from François et al. (Fig 4A, B). Both analyses revealed upregulation of inflammatory response pathways and p53 signaling (Fig 4A-C), consistent with previous findings showing increased numbers of reactive astrocytes, as well as expression of senescence-associated proteins in dysmorphic neurons and balloon cells.^17,18^ Notably, both datasets also showed a shared downregulation of OXPHOS-related pathways (Fig 4D), suggesting that defects in mitochondrial energy production are common features associated with mTORopathy.^19^

**Figure 4.**
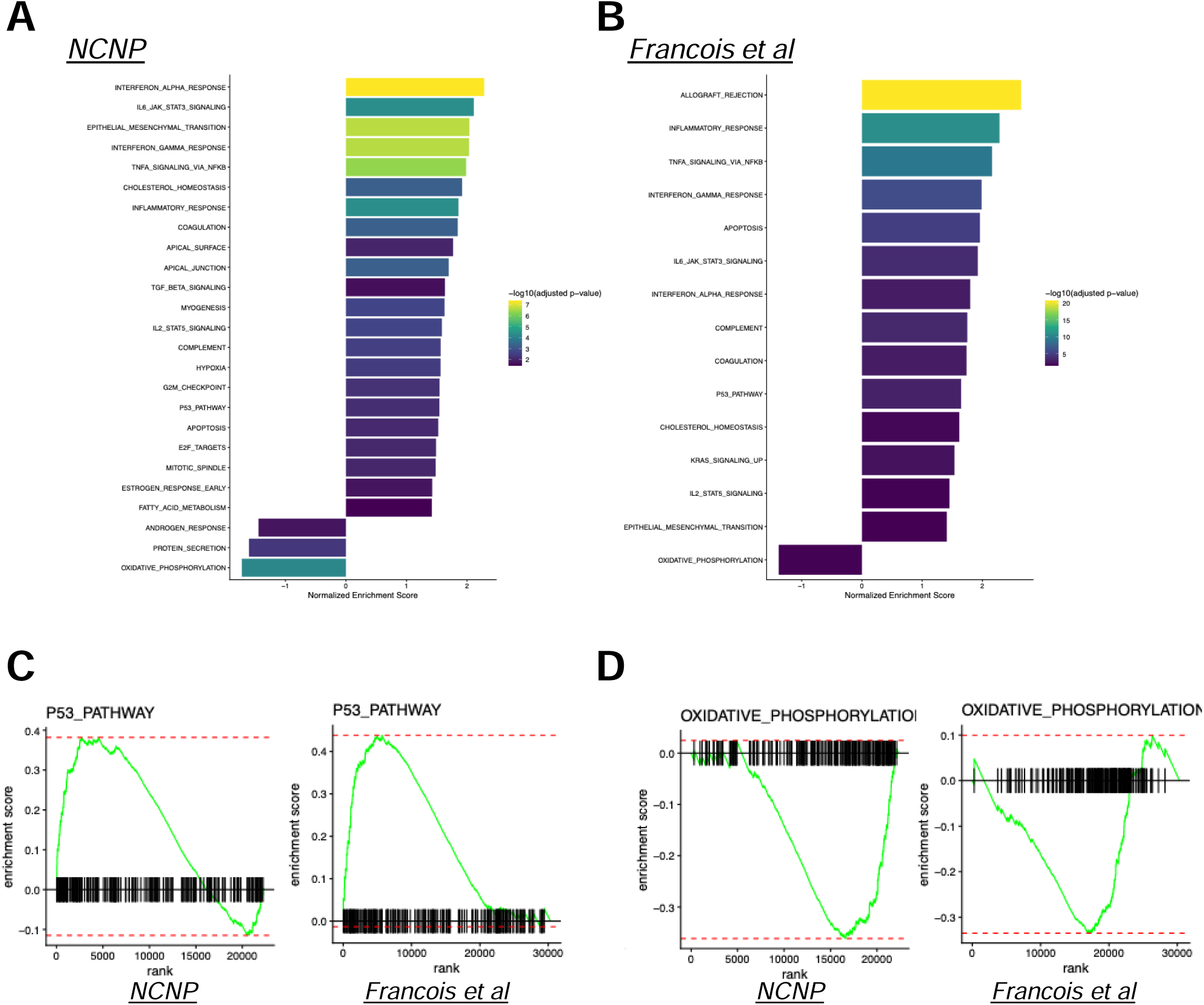
Functional enrichment analysis of DEGs. (A,B) Hallmark pathway enrichment analysis comparing NCNP and François et al. cohorts. (C, D) Gene set enrichment analysis (GSEA) plots for hallmark p53 signaling (C) and OXPHOS (D).

### Proteomic characterization of the mTORopathy brain

We next performed liquid chromatography–tandem mass spectrometry (LC–MS/MS) using surgically resected brain tissues obtained from control, MCD, and FCD III groups. To evaluate protein-level alterations specific to mTORopathy, we compared protein expression profiles between the control and mTORopathy groups. Expression levels of commonly detected proteins and genes showed a weak correlation (r = 0.29; Fig. 5A). Differential protein expression analysis identified 141 significantly upregulated and 223 significantly downregulated proteins in mTORopathy patients (Fig. 5B, Table S7). Notably, OXPHOS-related proteins, including COX5B,^20^ NDUFS4,^21^ GOT2,^22^ DKK3,^21^ and RAB27B^23^ were detected as differentially expressed proteins in this analysis (Fig. 5B). All of these OXPHOS-related proteins were significantly downregulated in the mTORopathy group, whereas their expression levels were not consistently altered in the FCD I or FCD III groups (Fig. 5C-G). Together, these findings suggested that the impairment of OXPHOS represents a specific molecular feature of mTORopathy brain tissue.

**Figure 5.**
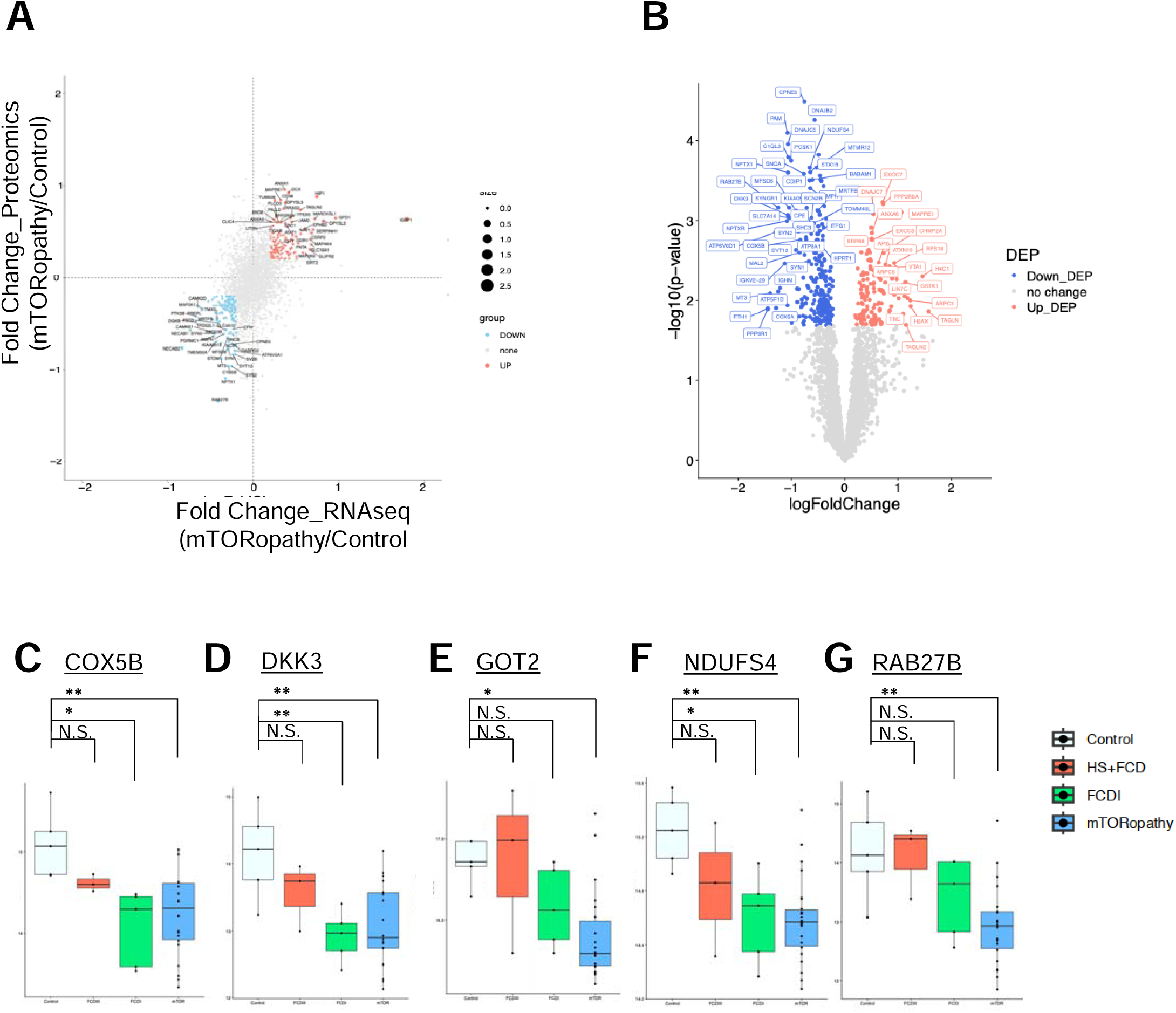
Integrated transcriptomic and proteomic analyses in mTORopathy patients. (A) Correlation of RNA-seq and proteomics fold changes (mTORopathy vs. control). Genes consistently altered at the transcript and protein levels are highlighted. (B) Volcano plot of differential expressed proteins (DEP) in mTORopathy samples. (C–G) Box plots of protein abundances decreased in mTORopathy: COX5B (C), DKK3 (D), GOT2 (E), NDUFS4(F), RAB27B (G). Statistical significance indicated: *\*P* ≤ 0.05, *\*\*P* ≤ 0.01, *\*\*\*P* ≤ 0.001., N.S. = not significant.

## Discussion

mTORopathy is a subgroup of malformations of cortical development (MCDs), encompassing cortical malformation disorders such as tuberous sclerosis complex (TSC), focal cortical dysplasia type IIa and b (FCD IIa/b), and hemimegalencephaly (HME). Recent transcriptomic studies using bulk RNA-seq and single-nucleus RNA-seq revealed alterations in pathways related to energy metabolism, synaptic plasticity, and inflammatory responses.^7,16,24,25^ However, protein-level analyses remain limited. Since PI3K/AKT/mTOR signaling is primarily regulated through protein-level networks, comprehensive proteomic profiling is critical for elucidating the pathology of mTOR-related epilepsy.

In this study, we performed integrated transcriptomic and proteomic analyses and provided comprehensive datasets as a bioresource for the research community. A unique feature of our dataset is that both transcriptomic and proteomic profiles were largely obtained from the same individuals. In total, 56 patients were enrolled, and 22 patients provided both RNA-seq and proteomic data. Although the number of proteins detected by proteomics analysis was smaller than the number of transcripts identified by RNA-seq, transcript and protein expression levels were positively correlated. Thus, these data enable integrative analysis across transcriptomic and protein levels and serve as a valuable resource for advancing the understanding of the molecular mechanisms underlying mTORopathy.

One candidate pathway regulating epileptogenesis in mTORopathy is the oxidative phosphorylation (OXPHOS) signaling pathway. GSEA analysis revealed significant downregulation of the OXPHOS signaling pathway at the transcriptomic level, as well as differential protein analysis identified reduced protein abundance of key OXPHOS proteins, including COX5B, RAB27B, NDUFS4, GOT2, and DKK3. Cytochrome c oxidase subunit 5B (COX5B) and NADH:ubiquinone oxidoreductase subunit S4 (NDUFS4) are mitochondrial proteins essential for the electron transport chain and ATP synthesis .^20,21,26,27^ Glutamate-oxaloacetate transaminase 2 (GOT2), localized in the inner mitochondrial membrane, functions in the malate–aspartate shuttle to transfer electrons from cytosolic NADH and thereby support ATP synthesis in mitochondria.^22^ Therefore, impairment of these protein functions drastically reduces OXPHOS signaling activity and ATP synthesis. Moreover, reduced expressions of RAB27B^23^ and Dickkopf-related protein 3 (DKK3) have been reported to decrease OXPHOS signaling activity.^28^ Together, these findings suggest that the dysregulation of key OXPHOS proteins may lead to impaired OXPHOS activity and diminished ATP synthesis in mTORopathy. Of note, OXA1L was identified as an upregulated protein in mTORopathy brain tissue. OXA1L is a mammalian homolog of Oxa1 in yeast^29^, facilitating the mitochondrial inner membrane insertion of OXPHOS proteins and promoting OXPHOS activity.^30^ Since our data revealed the reduction of COX5B and NDUFS4, core components of OXPHOS, OXA1L upregulation might not affect the OXPHOS activity profoundly.

As OXPHOS represents the principal source of cellular energy production critical for neuronal function,^31^ mitochondrial dysfunction, including impairment in OXPHOS activity, directly contributes to network hyperexcitability and seizure susceptibility.^32,33^ Consistent with our findings, François et al. reported dysregulation of energy metabolism, including OXPHOS signaling, as a conserved feature across epileptic conditions.^16^ FDG-PET studies have repeatedly demonstrated focal hypometabolism in focal cortical dysplasia,^14,15^ TSC,^34^ and HME,^35^ suggesting regionally reduced glucose uptake in mTORopathy. Reduced OXPHOS signaling may therefore provide a molecular basis for this hypometabolic phenotype in mTORopathies.

Mitochondrial dysfunction has also been observed in human epileptic brain tissues.^36,37^ Furthermore, inherited mutations in OXPHOS-related genes have been associated with familial epilepsy^38^, supporting the pathogenic role of OXPHOS signaling and highlighting it as a potential therapeutic target for future studies.

Our dataset was generated using bulk RNA sequencing and bulk proteomics, together with somatic mutation profiling. Because bulk tissue contains a mixture of neurons, glial cells, vascular cells, and blood-derived cells, the observed transcriptomic and proteomic alterations may reflect changes in both cellular composition and cell-type-specific gene regulation. Therefore, we could not conclude which cell types primarily drive the molecular signatures identified in this study. In addition, clinical variables such as age at surgery, sex and medication history differed across patients, which may have contributed to inter-individual variability, although genetic background heterogeneity was minimized because all samples were derived from a single-institution Japanese cohort. Future studies using larger independent cohorts and single-cell or spatially resolved multi-omics approaches will be required to refine the cellular origins of these molecular changes and to further elucidate the mechanisms of epileptogenesis in mTOR-related epilepsy.

In conclusion, our integrated transcriptomic and proteomic datasets, together with publicly available datasets, provide valuable insight into the molecular mechanisms underlying epileptogenesis and may contribute to the identification of therapeutic targets for drug-resistant epilepsies.

## Data availability

Raw sequence files obtained in this study will be available from DDBJ after publication. Before that, raw sequence files can be shared upon request.

## Acknowledgements

We greatly appreciate the patients and their families for participating this study. Illustrations in the figures were created with BioRender.

---

**Table 1:**

| Gene name | Mutation | Mutation type | ClinVar |
| --- | --- | --- | --- |
| *MTOR* | p.Thr518Ala | Ms | N.D. |
| *MTOR* | p.Leu1460Pro | Ms | Pathogenic |
| *MTOR* | p.Thr1977Lys | Ms | Pathogenic |
| *MTOR* | p.Ser2215Tyr | Ms | Pathogenic |
| *MTOR* | p.Ser2215Phe | Ms | Pathogenic |
| *MTOR* | p.Leu2427Pro | Ms | Pathogenic |
| *TSC2* | p.Ala31fs | Fs | N.D. |
| *AKT3* | p.Glu17Lys | Ms | Pathogenic |
| *AKT3* | p.Leu51His | Ms | N.D. |
| *AKT1S1* | p.Cys167fs | Fs | N.D. |
| *PIK3CA* | p.Glu545Lys | Ms | Pathogenic |
| *PIK3R2* | p.Pro4Ser | Ms | Benign |

Ms: Missense, Fs: Frame shift, N.D: No data

